# Illuminating microbial metabolic activities in the dark deep ocean with metaproteomics

**DOI:** 10.1101/310524

**Authors:** Zhang-Xian Xie, Shu-Feng Zhang, Hao Zhang, Ling-Fen Kong, Lin Lin, Da-Zhi Wang

## Abstract

The deep ocean is the largest habitat on earth and holds diverse microbial life forms. Significant advances have been made in microbial diversity and their genomic potential in the deep ocean, however, little is known about microbial metabolic activity that is crucial to regulate the bathypelagic carbon sequestration. Here, we characterized proteomes covering large particulate (>0.7 μm), small particulate (0.2-0.7 μm) and dissolved (10 kDa-0.2 μm) fractions collected at a depth of 3000 m in the South China Sea. The *Rhodospirillales,* SAR324, SAR11, *Nitrosinae/Tectomicrobia* were the major contributors in the particulate fraction whereas *Alteromonadales* and viruses dominated the dissolved counterpart. Frequent detection of transcription or translation proteins in the particulate fractions indicated active metabolism of SAR324, Archaea, SAR11, and possible viable surface microbes, e.g. *Prochlorococcus*. Transporters for diverse substrates were the most abundant functional groups, and numerous spectra of formate dehydrogenases and glycine betaine transporters unveiled the importance of methylated compounds for the survival of deep-sea microbes. Notably, abundant non-viral proteins, especially transporters and cytoplasmic proteins, were detected in the dissolved fraction, indicating their potential roles in nutrient scavenging and the stress response. Our size-based proteomic study implied the holistic microbial activity mostly acting on the labile dissolved organic matter as well as the potential activities of surface microbes and dissolved non-viral proteins in the deep ocean.

**Importance:** The deep ocean produces one third of the biological CO_2_ in the ocean. However, little is known about metabolic activity of the bathypelagic microbial community which is crucial for understanding the biogeochemical cycling of organic matter, especially the formation of bulk refractory dissolved organic matter (DOM), one of the largest reservoirs of reduced carbon on Earth. This study provided the protein evidence firstly including both particulate and dissolved fractions to comprehensively decipher the active microbes and metabolic processes involved in the DOM recycling in the deep ocean. Our data supported the hypothesis of the carbon and energy supply from the labile DOM after the solution of sinking particles to the bathypelagic microbial community.

## INTRODUCTION

The vast areas of deep ocean, characterized by low temperature, high hydrostatic pressure and complete darkness, is the most unexplored biome on Earth and produces one third of the biological CO_2_ in the ocean (1). However, this habitat has not yet been greatly explored. Recently culture independent sequencing techniques, i.e. 16S rRNA gene sequencing (2–7) and metagenomics (8–11), have been applied to explore the phylogenetic diversity and genomic potential of microbial communities in the deep ocean. Bacterial (5), archaeal (12) and small eukaryotic (3) populations are diverse and abundant in this extreme environment. Single-cell genomics provides the first insights into the unambiguous genome features of several uncultured clades from the deep ocean, i.e. SAR324, Arctic96BD-19, Agg47 and a deep SAR11 bathytype (13–15). Despite of this progress, very little is known about microbial metabolic activities in the deep ocean because of methodological limitations, but such knowledge is essential to understand the biogeochemical cycling of organic matter, especially the formation of bulk refractory dissolved organic matter (DOM), one of the largest reservoirs of reduced carbon on Earth (16).

Proteins as the carrier and the functional executor of life can indicate information on metabolic activities and provide clues into the decomposition and degradation of organic matter (17). Recently, the emergence of metaproteomics has greatly broadened our knowledge of marine microbial activities by identifying proteins in the particulate fraction (18–24) and the dissolved fraction (25–28). However, these studies typically focus on a certain microbial fraction, resulting in partial snapshots of *in situ* microbial behavior. In addition, most of these studies are conducted on the surface ocean: no systematic effort has been devoted to the proteins from the dark deep ocean.

As the depth increases in the ocean, the quantity and quality of DOM decrease (16). The microbial carbon pump (MCP) hypothesis suggests that the fresh produced DOM are rapidly removed by microbial metabolism in the surface and mesopelagic layers, leaving a background concentration of bathypelagic DOM refractory due to either chemical resistance or too dilute for microbial utilization (16). Although microbial activity in the deep ocean is key to determine the fate of DOM deposited there and to support the MCP hypothesis, understanding of the microbial activity is very lack. A recent metaproteomics of the free-living fraction (0.2-0.8 μm) indicates that the heterotrophic community utilize similar DOM from the lower euphotic and bathypelagic layers (29). However, this study mainly focuses on the transporters, and meanwhile, information of microbial activity beyond the size fraction is missing. Despite of that, it leads to our hypothesis that the microbial community in the deep ocean is more dependent on the labile DOM rather than refractory DOM. Here we applied a shotgun proteomic approach to explore the proteomes covering large particulate (LP, >0.7μm), small particulate (SP, 0.2-0.7 μm) and dissolved (DS, 10 kDa-0.2 μm) fractions collected from the dark deep ocean (3000 m) of the South China Sea (SCS). Using combined bacterial and viral metagenomic databases, 315 non-redundant proteins were confidently identified: they exhibited diverse microbial origins and functions, providing first insights into the holistic microbial metabolic activities in the dark deep ocean.

## RESULTS

### Proteomic overview

Four metagenomic datasets (8, 10, 30) (Table S1) were combined for protein identification in this study, including three bacterial metagenomics (the local DCM community, SEATS_DCM; the vertical community of Hawaii Ocean Time-series, HOT; and the Mediterranean Sea community at a depth of 3010 m, Deep_Med) together with a viromic dataset (Pacific Ocean Virome, POV, with size < 0.2 μm). Searching against the combined reference database, 228, 24 and 104 proteins matching 2112, 136 and 631 spectra were confidently identified from the LP, SP and DS fractions, respectively (Table S2). After removing redundant proteins among the three fractions, 315 proteins remained. The local surface metagenomic dataset (SEATS_DCM) obtained the most sequences in each fraction, followed by the vertical community genomics of HOT. The dataset of the Pacific Ocean Virome (POV), targeted at a size below 0.2 μm, presented hits comparable with the metagenomics of the deep Mediterranean Sea (Deep_Med) in the LP fraction, but displayed more hits in both SP and DS fractions. Notably, the protein profile of the DS fraction was composed almost half and half of sequences from SEATS_DCM and POV. Moreover, only 27 and 15 proteins matched the sequences from 4000 m of the HOT station in the Pacific Ocean and 3010 m of the Mediterranean Sea. Surprisingly, only 38% of the total 315 proteins matched the sequences from the dark ocean if 200 m down from the surface was thought of as the upper boundary of the dark ocean (Table S1).

### Biological origins

More than half of the spectra identified in each fraction originated from bacteria, followed by eukaryotes in the LP and SP fractions but by viruses in the DS fraction (Fig. 1A). Proteins in the LP fraction exhibited the most diverse microbial origins (Fig. 1A). *Rhodospirillales,* SAR324, *Nirospinae/ Tectomicrobia* and SAR11, and picophytoeukaryotic prasinophytes contributed the most spectra in the LP fraction. Most spectra from the archaea were detected only in the LP fraction, accounting for less than 5% and almost all of them from the thaumarchaea. Most of proteins in the SP fraction originated from *Alteromonadales, Enterobacterales, Nitrosinae/Tectomicrobia* and prasinophytes. Specifically, *Alteromonadales,* and prasinophytes dominated both proteomes of the SP and DS fractions. Other bacterial spectra from *Rhodobacterales,* SAR324, *Betaproteobacteria,* SAR11 and SAR116 were frequently detected in the DS fraction. Most of the viral spectra were found only in the DS fraction and many were assigned to cyanophages and uncultured phages from the DCM of the Mediterranean Sea. Five proteins representing 0.4% of the spectra in the LP fraction and 4.2% in the DS fraction were close to viral structural proteins but assigned to the bacteria; they might be from uncharacterized viruses or viral genes integrated to bacterial genome. Surprisingly, surface phytoplankton, i.e. *Prochlorococcus*, *Synechococcus* and prasinophytes, significantly contributed to the proteomes of all fractions, indicating that allogeneic microbial groups made up an important part of the microbial community in the dark deep ocean. It should be pointed out that quite a number of spectra in the LF (11%) and DS (28%) fractions were not assigned to any known organisms.

**Fig. 1.**
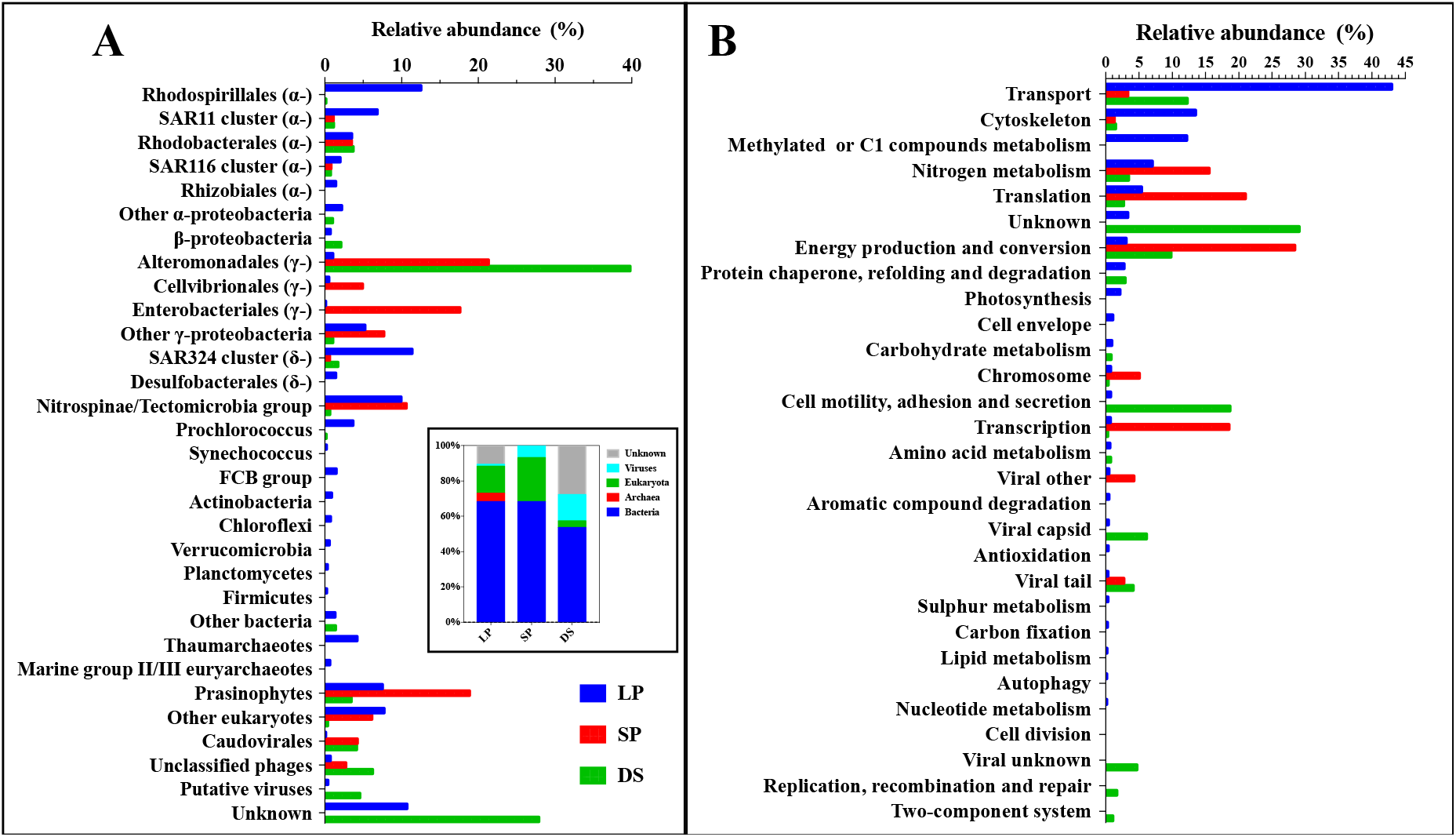
Taxonomic (A) and functional (B) distribution of proteins from the deep SCS. Percentages of each protein category show their contributions to proteomes in each fraction. The inset in Fig. 1A shows the distribution of the superkingdoms in each fraction. LP: large particulate fraction (>0.2 μm); SP: small particulate fraction (0.2 - 0.7 μm); DS: dissolved fraction (10 kDa - 0.2 μm).

### Protein functions

Protein function was interpreted using annotations based on the NCBInr, COGs, Pfam and KEGG databases (Fig. 1B). Except for the unknown functions, spectra assigned to transport, cytoskeleton, Methylated and one-carbon compounds metabolism, nitrogen metabolism, transcription and translation, energy production and conversion, protein chaperones, refolding and degradation, and photosynthesis were abundant in the LP fraction; while spectra affiliated with translation and transcription, energy production and conversion, and nitrogen metabolism dominated in the SP fraction. Many spectra assigned to viral structure proteins, i.e. capsid and tail, were detected in the DS fraction. In addition, spectra related to cell motility, adhesion and secretion, transport, energy production and conversion, nitrogen metabolism, transcription and translation, and protein chaperones, refolding and degradation were frequently detected in the DS fraction.

Transporters were abundant in the proteome of the deep SCS (Fig. 1B), and 93 and 21 transporters representing 43 and 12% of spectra were detected in the LP and DS fractions but only one in the SP fraction. Diverse substrates and microbial origins were predicted to be affiliated with these transporters (Fig. 2A). The predicted substrates of the transporters were amino acid, polyamine, glycine betaine, oligopeptide, urea, and sugar in the LP fraction; while transporters for amino acid were frequently detected in the DS fraction. Regardless of abundance, not all transporters present in the LP fraction were found in the DS fraction, such as those for substrates of glycine betaine, oligopeptide, urea, sugar, ammonium, and 2-aminoethylphosphonate. *Alphaproteobacteria* (SAR11, *Rhodobacteceae*, *Rhodospiralles* and SAR116) and *Deltaproteobacteria* (SAR324 and *Desulfobacterales)* contributed most of the transporter spectra in the two fractions. Moreover, transporters for different substrates were species specific. Multiple periplasmic substrate-binding proteins for sugar, oligopeptide, carboxylate, nitrate, sulfonate and taurine, and amino acid were highly expressed in the SAR324 group; while transporters with less diverse substrates were found in SAR11, and even less in other bacteria (Fig. 2). In the DS fraction, high abundance of the TonB-dependent receptor was detected in *Alteromonadales*. Interestingly, high-affinity Pi transporters (PstS) from both SAR11 and *Alteromonadales,* and one putative transporter for 2-aminoethylphosphonate from SAR324 were detected, indicating that some of these microbial groups might be subjected to phosphorus limitation in the deep ocean. Transporters for urea, taurine, ammonium and nitrate were identified, suggesting that these nitrogen-containing compounds were important nitrogen sources for the deep oceanic microbial community, although the community expressed abundant substrate-binding proteins for amino acid, oligopeptide and polyamine.

**Fig. 2.**
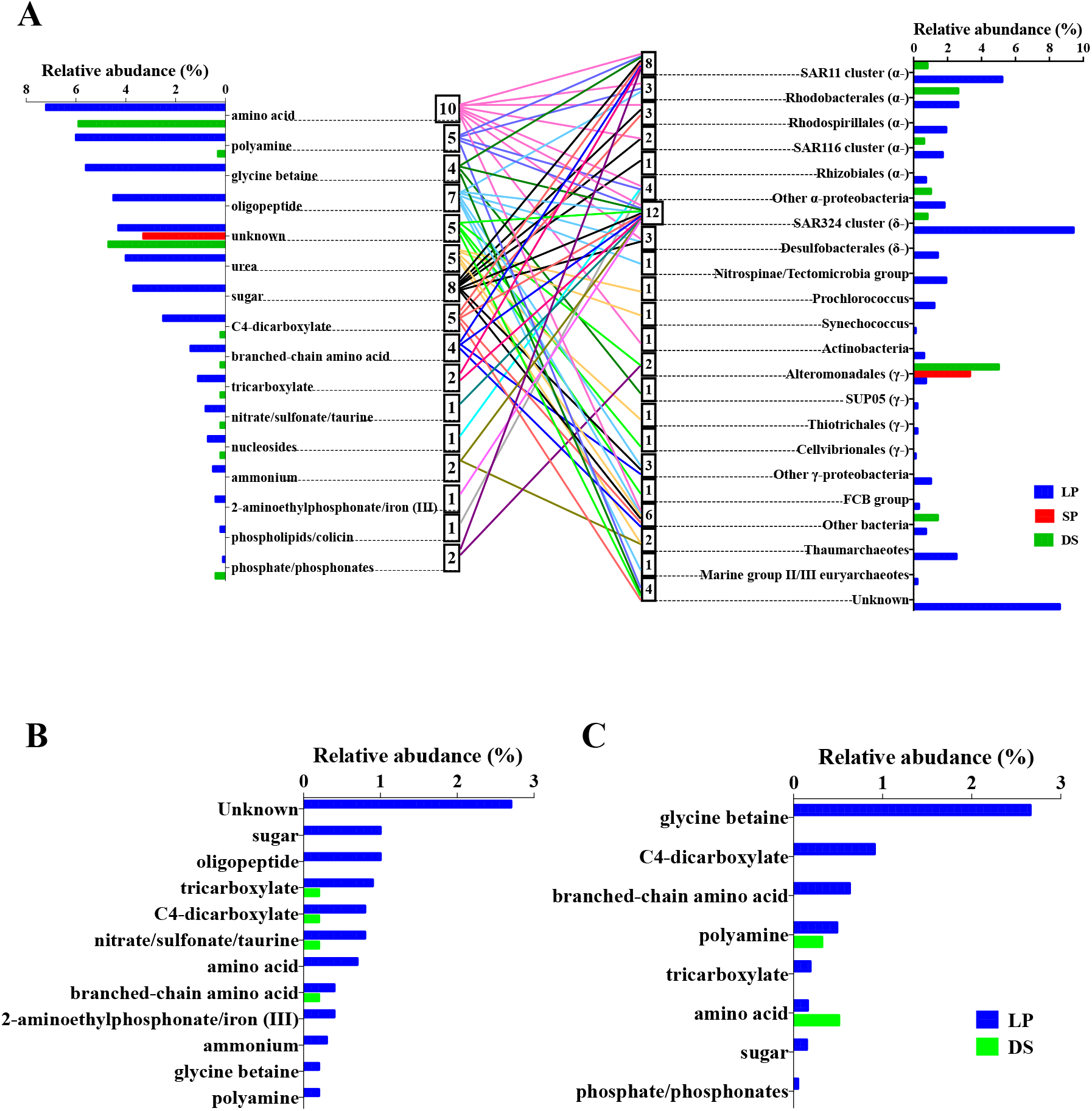
Relative abundance of transporters in each fraction in terms of predicted substrates and taxonomy (A), substrates of transporters from SAR324 (B) and from SAR11 (C). Squares in Fig. 2A shows the numbers of connecting predicted substrates (left) and their microbial groups (right).

Several proteins related to transcription and translation, such as DNA-directed RNA polymerase subunit B and C (RpoB and RpoC), elongation factor Tu (Tuf) and 1-alpha (EEF1A), and ribosomal proteins were frequently detected in the LP fraction. These proteins were linked to several microbial groups, such as the *Gammaproteobacteria,* SAR324, SAR11, thaumarchaea, *Chloroflexi, Planctomycetes, Prochlorococcus, Betaproteobacteria* and prasinophytes (Table S3 and Fig. 3), indicating active metabolisms of these microbial groups in the dark deep ocean. Interestingly, these proteins were also found in the DS fraction, such as RpoC, Tuf and ribosomal protein L2 from the *Alteromonadales,* as well as RpoC from SAR324 and Tuf from the *Chromatiales*.

**Fig. 3.**
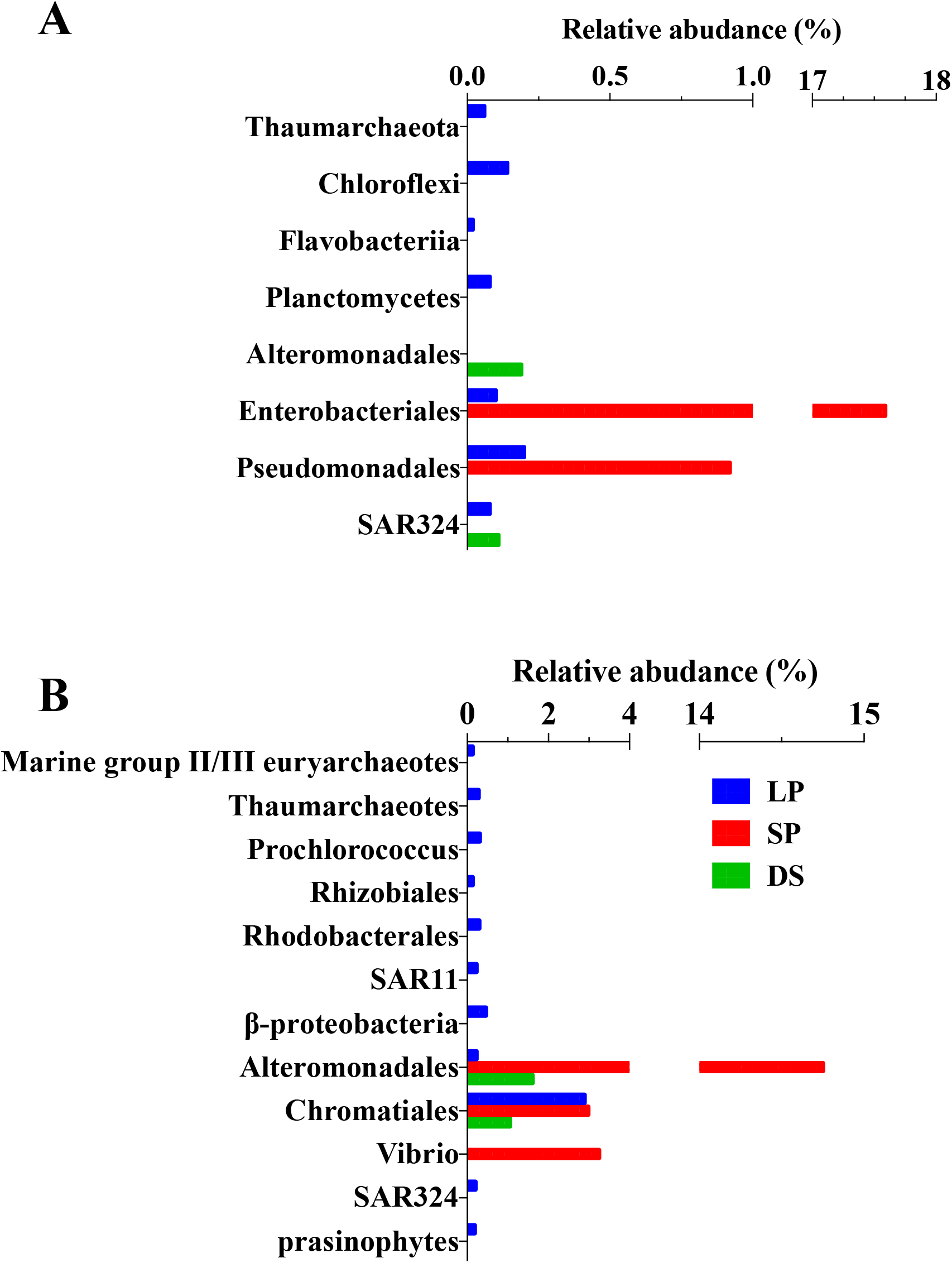
Relative abundance of RNA polymerase (A) and proteins related to translation (B) in terms of taxonomy.

The subcellular locations of all detected non-viral proteins were *in silico* predicted using CELLO (31), which apparently showed a fraction-specific pattern of cellular proteins in the deep ocean (Fig. 4). Cytoplasmic proteins contributed the most spectra in both LP and SP fractions but extracellular proteins instead in the DS fraction. Moreover, spectra assigned to cytoplasmic proteins, especially those from the *Alteromonadales*, were also abundant in the DS fraction, indicating that these proteins were protected from proteolysis.

**Fig. 4.**
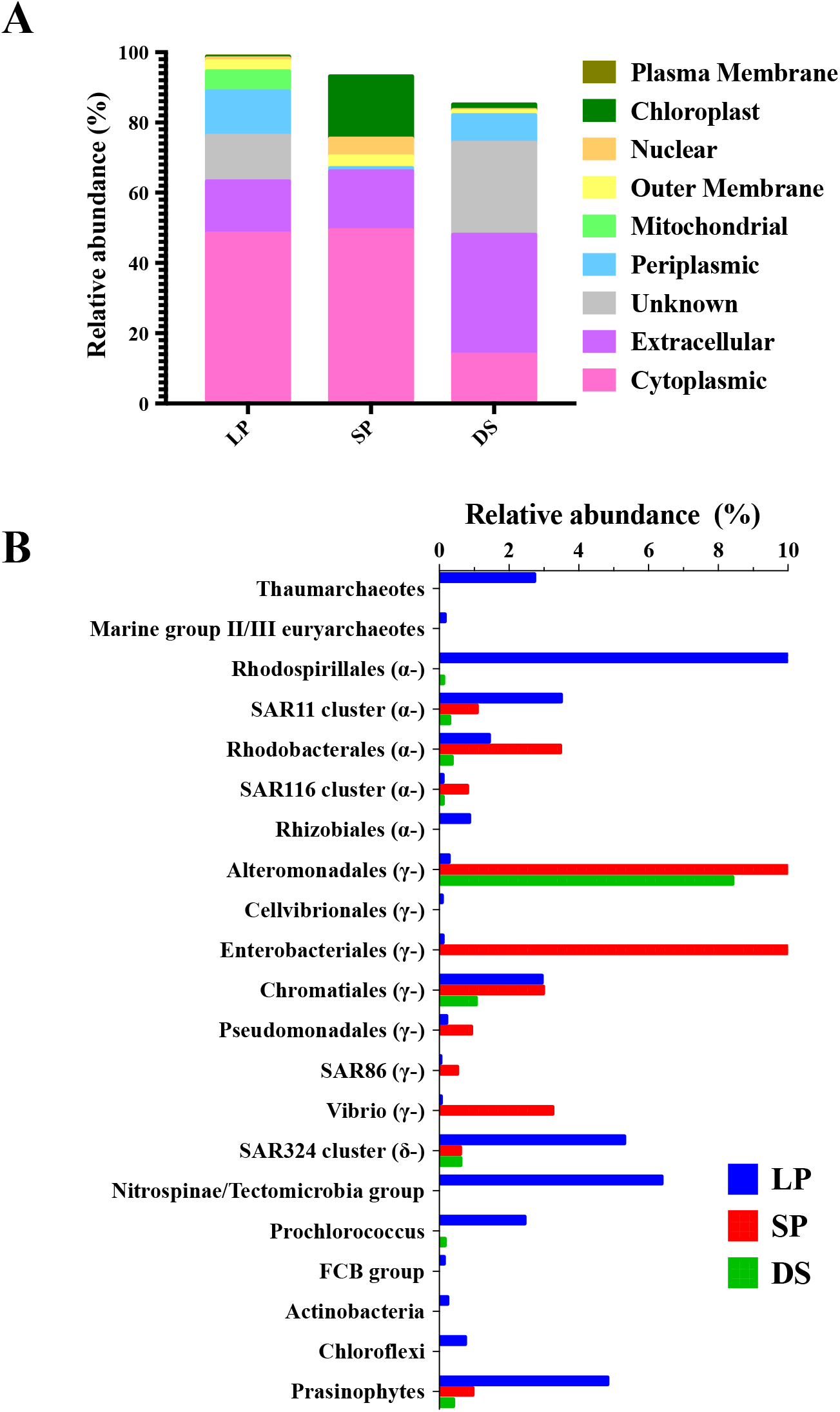
Relative abundance of proteins in terms of predicted subcellular location (A), and the taxonomic distributions of cytoplasmic proteins (B).

## DISCUSSION

Although great achievements have been made regarding the phylogenetic diversity and genomic potential of microbial communities in the dark deep ocean, our knowledge of microbial metabolic activity is far less understood. This study applied a shotgun proteomic approach to characterize the whole proteome of the microbial community from the deep ocean, and unveiled major microbial players and their metabolic activities in the dark deep ocean. High abundance of transporters was affiliated with diverse labile DOM, such as amino acid, polyamine, glycine betaine, peptide and sugar (Fig. 2A), which is consistent with the pattern found at the bathypelagic layers of the Atlantic Ocean (29). Labile and semi-labile DOM rapidly turn over in the upper layer, and leave a large amount of refractory aged DOM in the deep ocean (16). However, A recent study suggests that the deep ocean contains a large fraction of modern DOM derived from sinking particles (32). These data as well as the absence of proteins related to carbon fixation or metabolism of refractory DOM in our data support the hypothesis that the solution of sinking POM mostly contributed the carbon and energy source to the deep ocean microbes (29).

### Important microbial groups and their activities in the dark deep ocean

Spectra affiliated with transcription or translation indicated active microbial groups, including SAR324, archaea, SAR11 and *Alteromonadales* (Fig. 3). SAR324 contributed significant spectra in the proteome of the deep ocean (Fig. 1A), which was consistent with the current knowledge of SAR324 as a typically abundant bathypelagic group (7). Versatile metabolisms including carbon fixation, one-carbon metabolism, sulfur oxidation and alkane oxidation are genetically implicated (33). In our study, high abundant transporters affiliating with sugar, oligopeptide, carboxylate, nitrate, sulfonate and taurine, and amino acid were detected (Fig. 2B), suggesting that the heterotrophic lifestyle of SAR324 in the deep SCS depended on these labile organic compounds. It is reported that SAR324 genomic bins contain genes for degrading aromatic compounds with the byproduct of formate, as well as the formate dehydrogenase (FDH) gene (34). The detection of the FDH from SAR324 suggested that SAR324 could be fueled with formate, perhaps via this degradation process. The actyl-CoA synthetase (ACSS) and carbon monoxide dehydrogenase in the Wood-Ljungdahl pathway, or ribulose-1,5-biphosphate carboxylase-oxygenase in the Calvin-Benson-Bassham Cycle are proposed to be involved in carbon fixation of SAR324 (13, 34). In our study, we detected only ACSS from SAR324. Meanwhile, its close homologs in SAR324 genomes were clustered with genes involved in the pathway to produce acetate from compounds such as acetylated peptidoglycan or fucose (Fig. 5). It seemed that SAR324 in the deep SCS might express ACSS for acetate metabolism rather than carbon fixation. In addition, protein chaperonin of GroEL and DnaK, and a regulator of protease FtsH were found, implying that protein refolding and proteolysis were important for the survival of SAR324 in the harsh bathypelagic environment. Spectra of flagellin from SAR324 were also frequently detected, suggesting their particle-associated lifestyle (13). SAR324 might be one of the important players involved in the transformation between sinking POM and bathypelagic DOM.

**Fig. 5.**
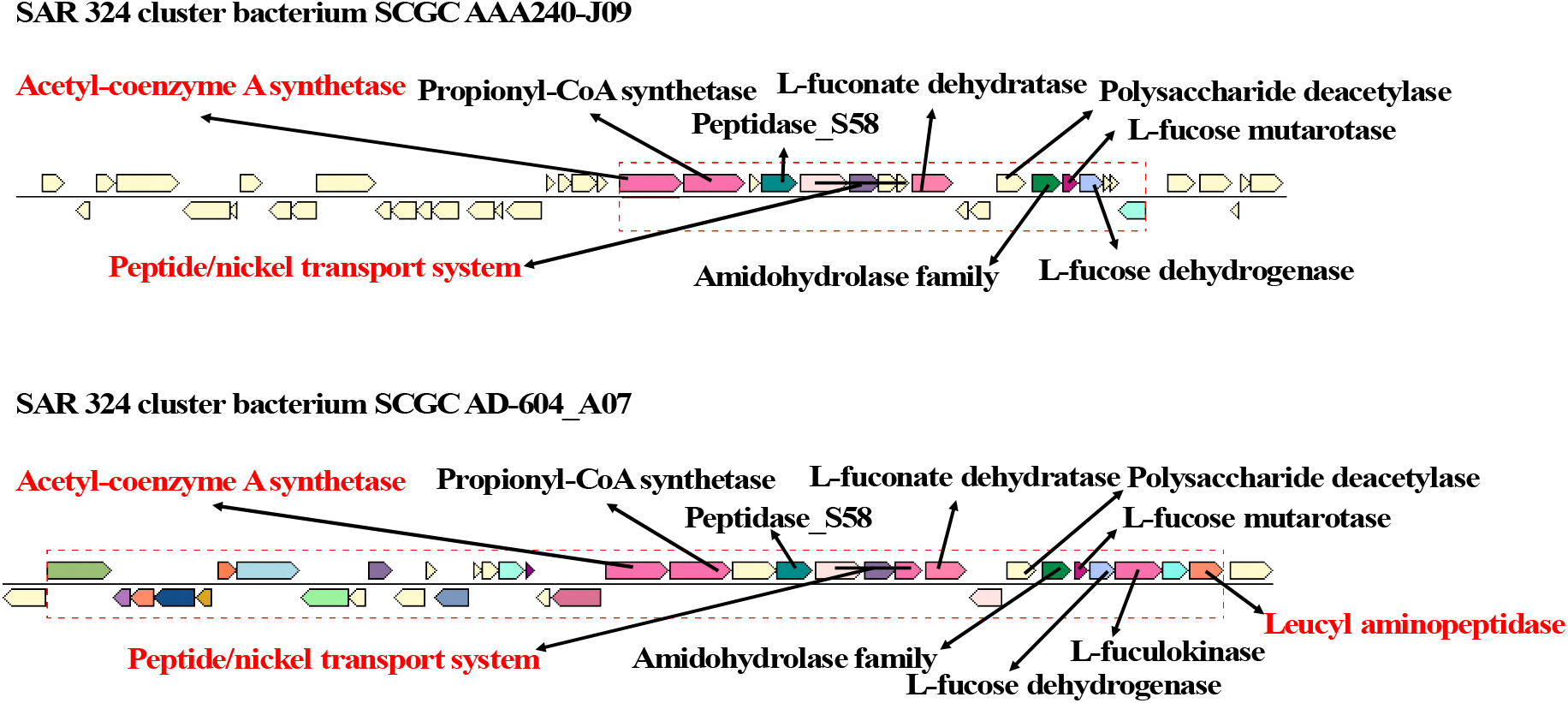
Genome context analysis of SAR324 contigs containing homologs of detected acetyl-coenzyme A synthetase, peptide/nickel system and leucyl aminopeptidase (in red). The detected ACSS of SAR324 in deep SCS appears to be related to acetate metabolism. The two gene clusters include both L-fuconate dehydratase and L-fucose dehydrogenase that are involved in the pathway of converting L-fucose to acetate. Nickel ion and oligopeptides e.g. acetylated peptidoglycan could be incorporated by SAR324 cells via peptide/nickel transport system. In the gene clusters, deacetylases such as N-acetylglucosamine-6-phosphate deacetylase in amidohydrolase family or polysaccharide deacetylase could catalyze peptidoglycan with the product of acetate. In addition, scavenging for nickel ion is required for the enzymatic activity of polysaccharide deacetylase.

SAR11 was another major contributor to the deep SCS proteome (Fig. 1A). Consistent with the dominance of transporters in the SAR11 metaproteome from the oligotrophic Sargasso Sea (20), spectra assigned to SAR11 in the deep SCS were also dominated by transporters, especially ABC transporters. A high proportion (13-16%) of transporter genes is found in the SAR11 genomes (35), but have distinct expression patterns in different systems. For example, amino acids, carboxylates, and polyamines are the dominant substrates of SAR11 transporters in the surface ocean (20, 22) whereas those for amino acid, sugar and taurine are abundant in a laboratory culture system (36). In our study, glycine betaine substituted for these compounds and became the preferred substrates in the SAR11 population from the deep SCS (Fig. 2C). In SAR11, methylated compounds such as glycine betaine is proposed to be tetrahydrofolate-mediated oxidized to formate with a final product of CO_2_ by FDH, and its utilization for energy production but not for biomass incorporation is experimentally demonstrated (37). The detection of FDH and the transporter profile in SAR11 suggested that the oxidation of methylated compounds including glycine betaine and formate were important energy sources for SAR11 to be a successful population in the energy limited deep ocean.

Cell division proteins FtsA and EEF1A from marine group II/III euryarchaeota, together with RpoB and EEF1A from the thaumarchaea, indicated that these two archaeal groups were active in the deep SCS, which was consistent with previous studies showing that they are abundant and active in the dark ocean (7, 12). Additionally, some other archaeal proteins were also detected. For example, pyruvate ferredoxin oxidoreductase (a key enzyme in the reverse Krebs cycle pathway for carbon fixation) was detected from marine group II/III euryarchaeota, indicating that the euryarchaeal autotrophic activity might occur in the deep SCS. Thaumarchaea in dark polar waters are reported to have urease genes that may enable them to utilize urea to fuel nitrification (38). Both detections of the urea transporter and ammonia permease in the thaumarchaea might supported the urea utilization by thaumarchaea in a cold and dark habitat for energy supply via nitrification.

Bacterial nitrification in the deep SCS was demonstrated by the detection of nitrite oxidoreductase from the nitrite oxidizing bacterium, *Nitrospina* (Table S3). Moreover, 5,10-methylenetetrahydromethanopterin-reductase (mer) involved in CO_2_ reduction to methane (39) indicated the presence of methanogenesis. Numerous spectra from FDH that was involved in energy generation during bacterial growth on C1 compounds (40) were detected (Table S3), demonstrating that formate might be an important DOM for many bacteria in the deep SCS.

### Surface microbial groups in the dark deep ocean

Proteins from cyanobacteria and prasinophytes were frequently detected, and the majority of them were found in the LP fraction (Table S4 and Fig. 1). It is common to capture phytoplanktonic proteins (18, 27, 41) and DNA (8, 10, 11) from the pelagic deep ocean, and sinking particles carrying surface phytoplanktonic detritus result in the detection of phytoplanktonic signals. However, a notable signal of translation related to *Procholorococcus* and *Bathycoccus prasinos* were observed in the proteome of the LP fraction, suggesting that these two phytoplanktonic groups might be active or in an infectious stage. Infected cells should be captured on the membrane. Proteins from *Synechococcus* phages rather than viruses infecting *Procholorococcus* or *B. prasinos* were detected in the LP fraction (Fig. S1A), even though *Synechococcus* cells were very fewer indicated by much less abundant *Synechococcus* proteins than the other two groups Fig. 1A), more likely supporting the active cells of *Procholorococcus* and *B. prasinos.*

Active and intact phytoplanktonic cells have been found at a depth of 500 m in the ocean (42–44) and even in the sea floor at a depth of up to 4500 m (45). So far, there is no phylogenetic evidence (42, 43) to support the presence of any native group of phytoplankton specifically occupying the habitats of the dark ocean, suggesting that the survived phytoplanktonic cells in the dark ocean are from the surface. Mechanisms for exporting surface phytoplankton to the deep ocean include sinking or fluffy particle association (43), internal solitary waves (42), or two-step transportation combining deep convection with down slope density current (44). However, it typically takes weeks for cells to reach this depth, hence phytoplanktonic cells face challenges to maintain their viability until they arrive at the deep ocean with extremely adverse environments. Probably, the picophytoplankton could survive with heterotrophic activity since several cyanobacterial strains have been demonstrated to have the ability to utilize organic compounds (46). The detection of FDH and ATP synthase from *Prochlorococcus* (Table S4) indicated that they might oxidize formate to CO_2_ for energy generation. Meanwhile, the finding of urea transporters and glutamate synthetase from *Prochlorococcus* indicated that it could use urea as a nitrogen source. Therefore, urea and formate might support *Prochlorococcus* to survive until cells reach the dark deep ocean.

Numerous spectra assigned to SAR11 were also detected in our study although the SAR11 of surface-ecotype and bathytype could not be distinguished from most of the peptides detected. Metagenomic recruitment suggests that surface SAR11 groups contribute more than 20% to the dark oceanic SAR11 population (15). Detection of a PstS protein in our study (Table S3) suggested that the surface SAR11 ecotype might contribute the population in the deep SCS, since that none of the single cell genome of the bathytype contains genes related to phosphate metabolism, i.e. PstS (15). Furthermore, the expression of SAR11 PstS is linked to phosphorus limitation in contrast to the relatively high concentration (3 μM) of phosphate at a depth of 3000 m in the SCS, which might be consistent with our suggestion.

Overall, our study suggested that the microbial community in the deep ocean might be comprised of live surface microbes and native microbes. The role of live surface microbes in the biogeochemical cycling of carbon, nitrogen and phosphorus in the deep ocean is an interesting concern.

### Viral and non-viral proteins in the DS fraction

Approximately 15% of the DS spectra were predicted to be virion-associated (Fig 1A) and many were related to cyanophages (Fig. S1A). Cyanophages were also found in the bacterial metagenomics of the dark ocean (8) as well as the proteomes of the LP and SP fractions in our study. These results suggested their contribution to the sinking carbons arriving in the deep ocean. Viral sequences from the dark ocean did not dominate the viral proteins (Fig. S2B), although the POV dataset included some deep-sea viromes (30). This might be partially caused by the lack of a local viromic database or less abundance of native deep-sea viruses. Numerous spectra originated from unclassified viruses, suggesting that the deep-sea viruses might be far less characterized. In the future, approaches such as integrated metaproteomics and metagenomics on purified environmental virions (47) combined with bioinformatic tools for environmental viral detection (48) could be applied to promote the characterization of unknown deep-sea viruses.

As in our previous studies (25, 27, 28), the DS samples without purification for virions allowed us to identify exoproteins referring to the proteins present in the extracellular milieu. After two-step filtration, cell density (< 10^3^ mL^−1^) of the DS fraction was below the detection limit of flow cytometry, indicating that the intact cells contributed little to the DS fraction. However, extracellular, cytoplasmic and periplasmic proteins were abundantly detected (Fig. 4). They can be derived from cell secretion, cell leakage or cell lysis, and some of them such as cytoplasmic protein may stay stable in marine extracellular milieu via being packed in nanogels or microgels (27) or having an enclosing membrane (49–51) to against enzymatic attack.

Exoproteins may indicate the interaction of cells with their environments (52). Except for functionally unknown (29%), proteins related to transport (12%) and motility (18%) dominated the DS fraction, which was consistent with both the *in silico* and experimental exoproteomes from marine bacteria (53, 54). Interestingly, the abundance patterns of transporters within a microbial group in terms of predicted substrates were sometimes not coincident between the DS and LP fractions (Fig. 2). For example, no spectra in the DS fraction were assigned to SAR324 transporters for sugar or oligopeptide despite of their relatively high abundance in the LP fraction, while low abundance taurine or branch-chain amino acid transporters from SAR324 were detected in both fractions. More obviously in SAR11, only transporters for polyamines and amino acids could be detected in the DS fraction, but both were less abundant. These results suggested that specific transporters in the DS fraction might be exported by different microbes, probably as function active exoproteins in the deep ocean.

Surprisingly, the spectral proportion of *Alteromonadales* (most was *Alteromonas macleodii*) was increased with the decreased size fraction, and reached the highest in the DS fractions (40%, Fig. 1A). Most of these spectra were associated with extracellular proteins linking to motility, transport and proteolysis. Interestingly, however, they also included typical cytoplasmic proteins, such as ribosomal protein L2, Tuf, glutamine synthetase and proteins related to the Krebs cycle (Fig. 4 and Table S5). The increasing cytoplasmic proteins found in the exoproteomes of marine bacteria in laboratory culture appears to be linked to cell lysis or leakage caused by adverse stresses (53). *A. macleodii* is a ubiquitous copiotrophic marine bacterium. The detection of flagellin was consistent the putative particle-attached lifestyle found in this group (55). As the attached sinking particles left the layer where they were living, the habitat became harsher: such as less available DOM and a cold temperature. Consequently, the ongoing cell lysis of *A. macleodii* probably occurred leading to only a few of their proteins being present in the larger fractions.

### Remark conclusions

Our proteomic study revealed the holistic metabolic activity of microbial community greatly dependent on the labile DOM in the dark deep ocean. Proteins for the uptake or oxidation of methylated compounds such as formate and glycine betaine were frequently detected, suggesting that these compounds might be an important carbon source for energy supply for the microbial community in the deep ocean. Both archaeal and bacterial nitrification also contributed energy production to the community. Microbial groups, such as SAR324, Archaea and SAR11 were the key metabolically active populations. Notably, the identification of translation proteins from phytoplankton such as *Prochlorococcus* indicated the presence of viable phytoplanktonic cells in the deep SCS. The picophytoplankton might survive via fueling by urea and formate, which raises the concern of the importance of surface microbes actively involved in the biogeochemical cycle of the deep ocean. In the DS fraction, abundant non-viral proteins, including cytoplasmic proteins and transporters, might indicate the physiological state of deep-sea microbial cells and their interactions with the harsh environments. These non-viral proteins should be ecologically important, since they might either be functionally active or served as an important labile or semi-labile DOM recycling in the ocean. In addition, the deep SCS proteome included numerous unknown viral sequences and taxonomically unclassified sequences, suggesting that much effort should be devoted to both viral and bacterial communities in the deep ocean.

## MATERIALS AND METHODS

### Sample collection

All three size fractions, each containing two replicates, were collected from the 3000 m depth at the SouthEast Asian time-series Study (SEATS) station (18.0 °N, 116.0 °E) in the SCS during the summer cruise of 2012. LP fraction samples (> 0.7μm) were achieved by filtering 1000 L seawater using an *in situ* large-volume water-transfer-system sampler (McLane) equipped with 142 mm GF/F glass-fiber filters (Whatman). A total of 110 L seawater for each replicate was collected with a rosette sampler equipped with 12-L Niskin Go-flo bottles (General Oceanics), and immediately filtered onboard through a GF/F glass filter. The materials captured on a subsequent Durapore membrane filter (0.2 μm, Millipore) were used for the SP fraction (0.2 - 0.7 μm), while the filtrates were collected as the DS fraction. The additional concentration steps for the DS fraction was were as previously described (28). In brief, the filtrates were concentrated onboard with a Millipore Pellicon 2 tangential flow filtration system with a 10 kDa regenerated cellulose filter, and further concentrated and desalted in a Millipore 50 mL stirred ultrafiltration cell on ice. All the membrane filters and concentrates were stored at −80 °C before protein extraction.

### Metaproteomic procedure

The procedure for protein extraction of the three fractions was slightly modified from our previous studies (18, 27, 28, 56). Briefly, filters for LP or SP fractions were cut into chips and suspended in lysis buffer (urea, thiourea, Triton X-100, carrier ampholytes,3-[(3-Cholamidopropyl)dimethylammonium]-1-propanesulfonate, protease inhibitor cocktail, and dithiothretol (DTT)), broken-down with an ultrasonic shaker, sonicated on ice, and finally centrifuged at 10000×g for one hour after one-hour incubation. The supernatant was transferred to a new tube while the sediment was rinsed with lysis buffer twice. All the supernatant was pooled together to be precipitated with ice-cold 20% trichloroacetic acid in acetone overnight. Protein extraction from the concentrated DS fraction was the same as in a previous study (28).

The protein extract was reduced with DTT and alkylated with iodoacetamide. Peptide solutions of the two replicates in each fraction were pooled after trypsin digestion on a 10 kDa Microcon filtration device (Millipore) using a filter-aided sample preparation method (57). The pooled peptide solution was separated with strong cationic exchange and desalted, then offline subjected to nanoelectrospray ionization connecting to a tandem mass spectrometry (Thermo Scientific Q Exactive™ Hybrid Quadrupole-Orbitrap™ Mass Spectrometer) as previously described (28). A data-dependent acquisition mode was applied to automatically select the 15 most abundant precursor ions for fragmentation by a high-energy collision induced dissociation with a dynamic exclusion duration of 15 s. A resolution of 70000 was used to MS scans from 350 Da to 2000 Da in the Orbitrap while a resolution of 17500 for fragmented ions. Tools in Proteome Discoverer (ver. 1.3.0.339; Thermo Fisher Scientific) were used to merge the raw data and transform to MGF format. The protein identification was performed using Mascot searching engine (ver. 2.3.0; Matrix Science, London, United Kingdom) as follows: specifying trypsin peptides and allowing one missed cleavage; 20 ppm and 0.6 Da mass tolerances for precursor and fragmented ions; fixed carbamidomethyl modification on cysteine and variable oxidation modification on methionine; peptide charges were set to +1, +2 and +3.

A custom reference database included four metagenomic datasets that are a local and contemporary metagenomics at DCM (SEATS_DCM, 75 m) (28); a suite of vertical community genomics from 10 m to 4000 m at the HOT station ALOHA (8); a metagenomics of the Mediterranean Sea at the 3010 m depth (Deep_Med) (10); and the largest dataset of viromics from 10 m to 4300 m in the Pacific Ocean (POV) (30). After removal of the redundant sequences, the reference database finally consisted of 7199369 sequences. Moreover, a reverse decoy database was automatically generated to test the proportion of false positive hits. Peptide identifications were accepted with a minimal probability of 95%. Mascot searches with a false discovery rate over 1% were rejected. Proteins matched with two or more peptides were finally accepted as confident identifications. The formation of “Protein Groups” was conducted and the highest scoring protein in the group was used as a master protein.

### Bioinformatic analysis

Taxonomic and functional annotation as well as subcellular location prediction for identified sequences were performed as previously described (28). To avoid the mis-annotation between virus and non-virus due to the auxiliary metabolic genes encoded by viruses, the BLASTp-based non-viral sequences from datasets of SEATS_DCM, HOT and POV (Deep_Med was ignored owing to no assembly data being available) were considered as putative viral sequences once they were annotated from viral contigs scanned using VirSorter (48). In addition, non-viral sequences functionally assigned to viral structure were also re-classified as putative viral proteins. Normalized spectral counts were used for proteomic semi-quantification following the previous description (47).

### Accession number(s)

The mass spectrometry proteomics data have been deposited in the Proteome-Xchange Consortium (58) (http://www.proteomexchange.org/) via the PRIDE partner repository with the data set identifier PXD00XXXX.

## ACKNOWLEDGEMENTS

We thank the captain and crew of the R/V DongFangHong 2 for their assistance. Professor John Hodgkiss is thanked for his help with English. Professor Matthew B. Sullivan is thanked for kindly providing the protein datasets of POV.

## Supplementary materials

**Fig. S1**. Percentages of viral spectra in proteomes of each fraction in terms of viral host (A) and sequence source (B).

**Table S1**. The numbers show how many sequences are identified in each dataset from the different fractions collected from the deep SCS (3000 m).

**Table S2** (see Excel file). Summary table of proteins identified from the deep SCS.

**Table S3** (see Excel file). Important proteins mentioned in the main text.

**Table S4** (see Excel file). List of identified proteins from surface phytoplankton.

**Table S5** (see Excel file). List of identified proteins from *Alteromonadales*.

